# Stabilizing Obligatory Non-native Intermediates Along Co-transcriptional Folding Trajectories of SRP RNA Affects Cell Viability

**DOI:** 10.1101/378745

**Authors:** Shingo Fukuda, Shannon Yan, Yusuke Komi, Mingxuan Sun, Ronen Gabizon, Carlos Bustamante

## Abstract

Signal recognition particle (SRP) in *Escherichia coli* comprises protein Ffh and SRP RNA. Its essential functionality—co-translational protein-targeting/delivery to cellular membranes— hinges on the RNA attaining a native long-hairpin fold that facilitates protein conformational rearrangements within the SRP complex. Since RNA folds co-transcriptionally on RNA polymerase, we use high-resolution optical tweezers to first characterize the mechanical unfolding/refolding of incrementally lengthened RNAs from stalled transcription complexes until reaching the full-length transcript. This analysis allows identification of folding intermediates adopted during the real-time co-transcriptional folding of SRP RNA. The co-transcriptional folding trajectories are surprisingly invariant to transcription rates, and involve formation of an obligatory non-native hairpin intermediate that eventually resolves into the native fold. SRP RNA variants designed to stabilize this non-native intermediate—likely sequestering the SRP ribonucleoprotein complex in an inactive form—greatly reduce cell viability, indicating that the same co-transcriptional folding mechanism operates *in vivo* and possible alternative antibiotic strategies.

**Highlights:**

1. Folding pathway of an essential functional RNA has been resolved co-transcriptionally.
2. The co-transcriptional folding pathway of SRP RNA is invariant to transcription rates.
3. Nascent SRP RNA obligatorily forms a non-native intermediate before adopting the native fold.
4. Modulating transitions from the non-native to native SRP RNA hairpin fold alters cell viability.

## INTRODUCTION

Nascent RNA transcripts, like their protein counterparts, begin to fold during their synthesis and before the entire sequence is completed and released from RNA polymerase (RNAP) (Al-Hashimi and Walter, 2008; Pan and Sosnick, 2006; Perales and Bentley, 2009). Accordingly, it has been speculated that the co-transcriptional folding pathway of RNA could differ significantly from that of the isolated full-length molecule, possibly comprising alternative folding intermediates. Differences in folding pathways hence may be particularly significant for functional RNAs, since their essential cellular activity hinges tightly on the attainment of specific folds. The mechanism underlying co-transcriptional folding of RNA has been investigated intensely by means of biochemical bulk assays (Chao et al., 1995; Heilman-Miller and Woodson, 2003; Lewicki et al., 1993; Nechooshtan et al., 2014; Palangat et al., 1998; Pan et al., 1999; Wong et al., 2005, 2007; Zhang and Landick, 2016) as well as computational methods (Zhao et al., 2011). Single molecule force spectroscopy has also been employed to monitor the folding of a *pbuE* adenine riboswitch, and its conformational switching upon ligand binding (Frieda and Block, 2012). However, a comprehensive analysis of the co-transcriptional folding/misfolding of an essential functional RNA at the single-molecule level, and its consequent biological implications, has not been described. For instance, does the vectorial (5’ to 3’) nature of RNA synthesis favor the folding and sequestering of the upstream RNA sequences into non-native intermediate structures? Are these 5’-end-sequestering intermediates devised to restore long-range base-pairing in the final functional RNA fold? Or are they in-route to misfold? Are *on-pathway* folding intermediates always *productive*, i.e. part of the final native state; or, alternatively, can a *non-productive* intermediate be *on-pathway*—i.e. obligatory—co-transcriptionally? Does the discontinuous nature of transcription—whereby RNA synthesis is interrupted by RNAP pausing (Artsimovitch and Landick, 2000; Larson et al., 2014; Gabizon et al., 2018)—play a role in determining the folding pathway? How does the rate at which the nascent RNA emerges from RNAP alter the folding process? As a functional RNA transcript emerges from RNAP, do changes in the population and residence time of the molecule in intermediate states lead to any biological consequences?

The 114-nucleotide (nt), highly conserved 4.5S RNA from *Escherichia coli (E. coli)* is a suitable model system to study the co-transcriptional folding of RNA. 4.5S RNA forms the core of the signal recognition particle (SRP) and is therefore referred to as SRP RNA (Doudna and Batey, 2004; Keenan et al., 2001). SRP RNA is known to adopt a simple, ~50-base-pair (bp) long hairpin functional form (Figure 1). It is in this native fold that SRP RNA orchestrates elaborate and critical conformational rearrangements, which serve as fidelity checkpoints for SRP to accurately target specific ribosome cargos and deliver them to appropriate translocation machineries in the membrane (Akopian et al., 2013). Specifically, SRP RNA recruits SRP protein FfH (Figure 1A, colored in blue) with pico-molar affinity. The resulting riboprotein complex forms the indispensable protein biogenesis SRP machinery, which targets ribosome cargos displaying specific nascent peptide chains. In *E. coli*, the G-domain of FfH (G for GTPase) must interact with the G-domain of the SRP receptor protein FtsY in the membrane (Figure 1A, colored in green). When FfH and FtsY first interact via their G-domains, the two proteins hang around the SRP RNA tetra-loop next to the hairpin *tip-proximal stem* (Figure 1A, cargo loading), where the FfH M-domain binds (Figure 1B, blue-shaded region). Next, the G-domains of the two proteins swing together from the hairpin tip, and dock onto the hairpin *tip-distal stem* of the RNA, (Figure 1A, activated SRP assembly) which comprises nucleotides 10-15 and 96-101. This docking configuration then triggers a tight binding of the G-domains and subsequently a close conformation that activates their GTP hydrolysis to signal cargo unloading (Figure 1A). The significance of the tip-distal stem of SRP RNA has thus been examined in great depth (Heilman-Miller and Woodson, 2003; Jagath et al., 2001; Peterson and Phillips, 2008; Shen et al., 2013; Siu et al., 2007). When SRP RNA’s distal stem is shortened by more than 14 bp (indicated by purple dashed line in Figure 1B), the docking configuration is impaired, which attenuates GTPase activity of the SRP?FtsY complex and abolishes ribosome cargo transport (Ataide et al., 2011). In general, the attainment of the SRP RNA native fold is essential for SRP functionality.

**Figure 1.**
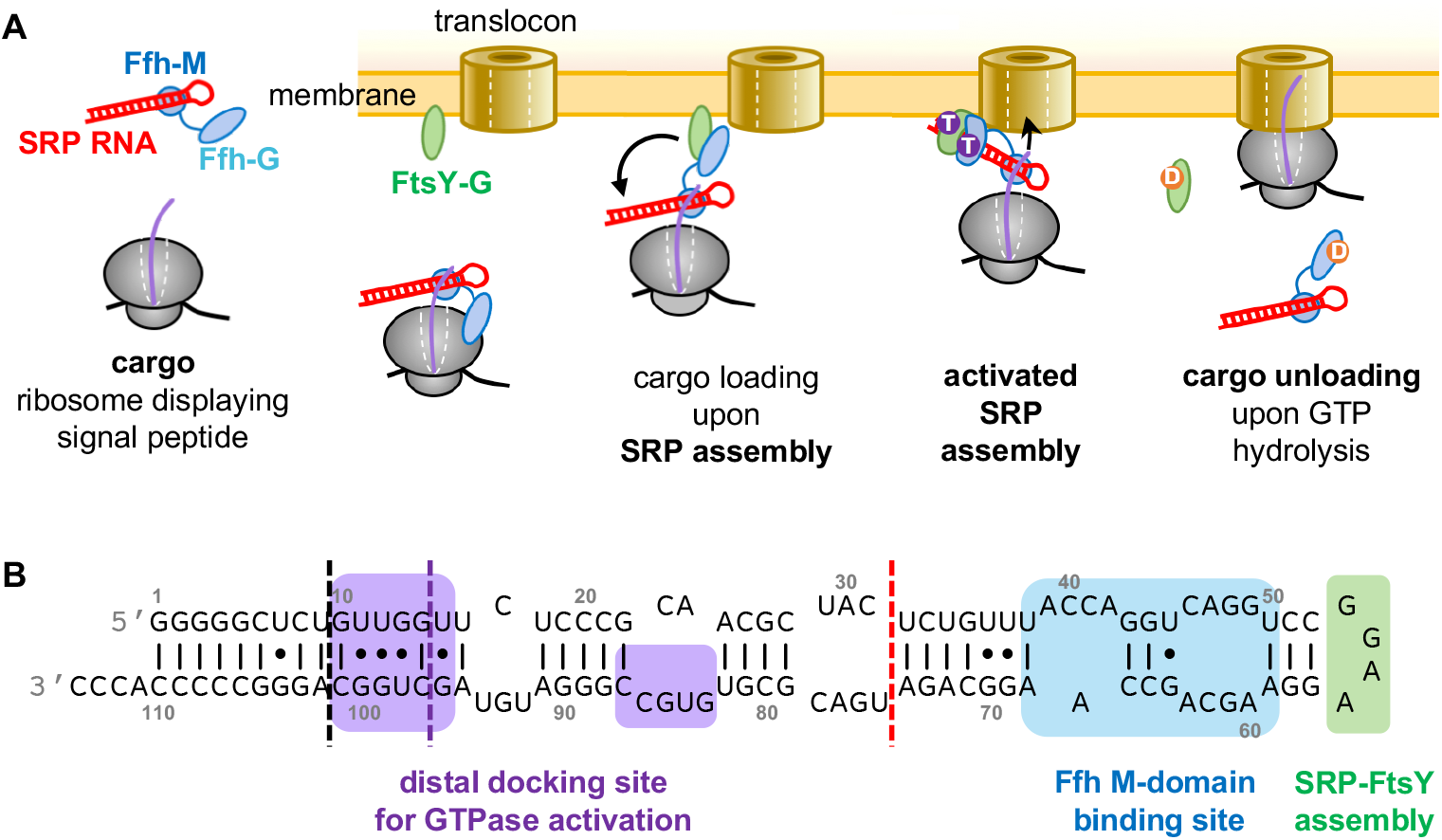
SRP protein-targeting activity. (A) Left-most: In *E. coli* the signal recognition particle, SRP, is composed of SRP RNA (hairpin fold in red) and SRP protein, Ffh (blue), where FfH M-domain binds to SRP RNA hairpin-tip proximal stem. Second-from-the-left: SRP then binds to ribosome cargos (grey) that display signaling peptide chains (purple), and brings the cargos to the vicinity of membrane-embedded translocons (brown barrels) by interacting with the membrane-bound SRP receptor protein, FtsY (green). This interaction is between the G-domains of Ffh and FtsY (Ffh-G and FtsY-G). Middle: The docking of both G-domains to the SRP RNA hairpin-tip distal stem triggers GTP hydrolysis on the two proteins (cartoon second-from-the-right), and activates cargo unloading from the SRP assembly to the translocon—through which the growing peptide chains will be subsequently transported across (right-most cartoon). (B) The native, functional long-hairpin fold of 114-nt long SRP RNA from *E. coli*. The hairpin-tip proximal stem where FfH M-domain binds is highlighted in blue; the hairpin tetra-loop region where the SRP-FtsY assembly initially interacts in green; and on the hairpin-tip distal stem highlighted in purple is the G-domain docking site, which is essential for SRP protein-targeting activity. Blunt-ended truncations on the long hairpin fold of SRP RNA (shown by colored dash lines) incrementally lower SRP GTPase activity, and eventually impair SRP protein-targeting functionality: black, no impact; purple, moderately lowered GTPase activity; red, GTPase activity completely abolished.

Here we use high-resolution optical tweezers (Comstock et al., 2011; Righini et al., 2018) to characterize the co-transcriptional folding pathway of individual SRP RNA molecules. To this end, we first characterize the structures adopted by incomplete nascent SRP RNA transcripts of various lengths emerging from *E. coli* RNAP molecules stalled at specific locations on the DNA template. Having established the folding signatures along the progression of a growing transcript, we then follow in real-time the emergence and co-transcriptional folding of SRP RNA on the surface of RNAP. We show that the folding trajectory of SRP RNA comprises regions of continuous transcript growth, sudden folding, concurrent growth and folding, and regions of transcriptional pauses. These details of the trajectory are remarkably robust from molecule to molecule and invariant to large changes in transcription rate, possibly reflecting the simple topology of the final native state. We observe the formation of a non-native, obligatory intermediate and its resolution into the final native state. Finally, we use the insights gained from uncovering this co-transcriptional folding pathway, to rationally engineer RNA mutations designed to alter specific structural folding transitions and characterize their impact on SRP’s cargo-targeting activity in *E. coli in-vivo*.

## RESULTS

### Characterization of SRP RNA Folding Intermediates from Stalled RNAP Complexes

To characterize intermediate folding structures of SRP RNA generated at different time points during transcription, we measured force-extension (F-X) curves of five truncated and the full-length SRP RNA using a high-resolution, home-built dual-trap optical tweezers instrument (Figure 2A; see Methods). We prepared a library of *E. coli* RNAP-RNA nascent chain complexes from DNA templates truncated at 40, 52, 84, 97, 114+16 and 114+39 bp, and stalled by a biotin:streptavidin roadblock at the DNA 3’-end. As the footprint size of RNAP is ~27-bp on the DNA and ~14 nt on the RNA (Komissarova and Kashlev, 1998; Monforte et al., 1990), the lengths of emerging RNA transcripts corresponding to the stalled complexes should be approximately ~13, 25, 57, 70, 103 and 114+8 nt, respectively (e.g. 40 - 14 - 27/2 = ~ 13 nt; complex I-VI in Figure 2B).

**Figure 2.**
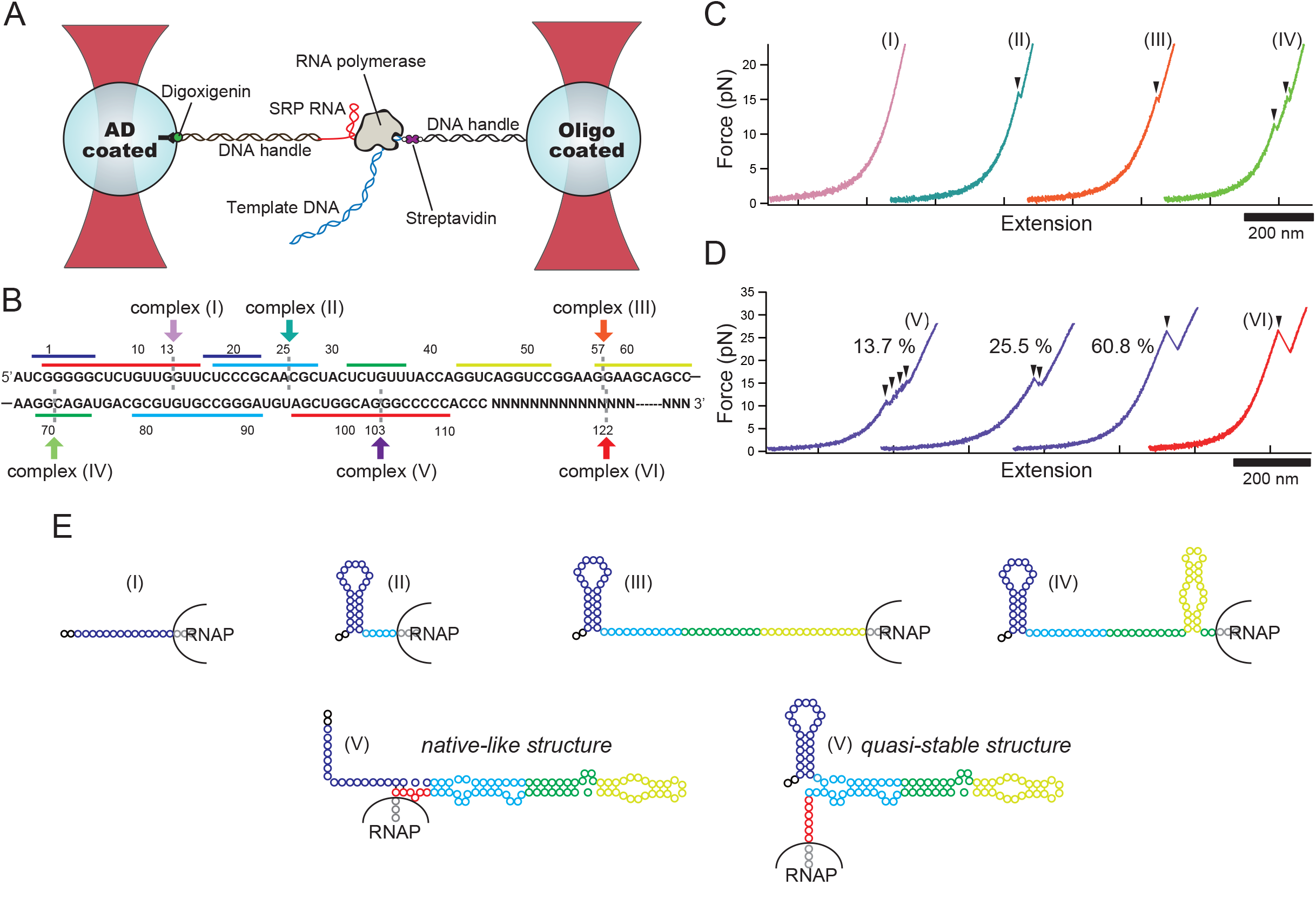
SRP RNA Sequence and Representative Force Extension Curves with Six Constructs. (A) Illustration of experimental setup to observe the force-extension curve of full-, truncated-SRP RNA structure with optical tweezers (not to scale). Two optical traps (red) were used to trap anti-digoxigenin (AD) and oligo coated beads. One DNA handle is attached to streptavidin for the roadblock of RNAP and the other handle is hybridized to nascent RNA chain. (B) SRP RNA sequence, including the color bars correspond to base-pairing when the form of native long hairpin structure. The blue bars show base-pairing when non-native intermediate structure forms. We prepared six DNA templates and these are truncated at (I) 40-, (II) 52-, (III) 84-, (IV) 97-, (V) +16- and (VI) +39-bp, respectively. For the constructs (V) and (VI), the DNA template were truncated at spacer region, where +1 corresponding to the position of incoming nucleotide followed by last nucleotide of SRP RNA sequence. Considering that RNAP covers ~27 bp on the template DNA and ~14-nt on the nascent RNA for the footprint, respectively, the lengths of RNAs emerging from RNAP are estimated to (I) ~13-, (II) ~25-, (III) ~57-, (IV) ~70- and (V) ~103-nt, respectively as shown by colored dashed lines. For the construct (VI), ~12-nt is flanked by full SRP RNA at 3’ end, allowing sufficient separation from RNAP surface. (C and D) Representative unfolding curves of truncated and full SRP RNA bound with RNAP: (I) ~13 (pink), (II) ~25- (light green), (III) ~57- (orange), (IV) ~70- (yellow green), (V) ~103-nt (purples) and (VI) full-SRP RNA (red), respectively. Arrow heads indicate unfolding events. (E) Expected secondary structures of truncated SRP RNA suggested by F-X curves

First, a force-ramp protocol was applied to repeatedly unfold and refold the RNA structures emerging on the RNAP surface. The resultant F-X curves show a number of rips whose size reflect the number of base-pairs of the transcript that unwound in each protocol. When we stretch the nascent chain complex I in Figure 2B, i.e. ~13-nt outside of RNAP, we do not observe any unfolding signature (pink curve in Figure 2C; complex I in Figure 2E). When SRP RNA is allowed to grow further (~25- and 57-nt long, complex II and III in Figure 2B), we observe characteristic transitions in the F-X curves (teal and orange curves in Figure 2C; complex II and III in Figure 2E), where the nascent RNA unfolds in a single rip at a mean force (F_unf_) of 15.4 ± 1.1 pN (Figure S1A). Using the wormlike chain model (Bustamante et al., 1994), we calculate an increase in contour length (ΔL_c_) of 12.8 ± 1.4 nm upon nascent chain unfolding (Figure S1B). This length increase is consistent with unfolding of a *non-native intermediate structure* (23-nt × 0.59 nm/nt - 2.2 nm/hairpin helix width = 11.37 nm), which was proposed previously in bulk studies (Watters et al., 2016; Wong et al., 2007). It was speculated that, during early transcription, this intermediate structure sequesters the 5’ upstream portion of SRP RNA by adopting a labile hairpin fold with a ~6-bp stem and an 11-nt loop. When the nascent RNA is ~70 nt long (complex IV in Figure 2B), its F-X curve shows two unfolding rips (light green curve in Figure 2C) indicating that—in addition to the first non-native intermediate—a second intermediate hairpin has formed, likely involving the native hairpin tip-proximal stem (Figure 2E, yellow hairpin in complex IV), consistent with structural predictions by Mfold (Zuker, 2003). When the transcript reaches ~103-nt in length (complex V in Figure 2B), we observed three types of F-X curves (purple curves in Figure 2D, N=153 unfolding events). The majority of F-X curves show one single large rip (60.8 %) and their shape is comparable to that of full-length SRP RNA (discussed below), indicating the formation of a *native-like structure* (Figure 2E, bottom left). The other two types of F-X curves display either two consecutive rips (25.5%) or a series of small rips reflecting gradual unfolding (13.7%), respectively; these features suggest the formation of *quasi-stable structures* involving the non-native hairpin intermediate (Figure 2E, bottom right). During repeated pulling and relaxation cycles, the F-X curves of complex V interconvert among these alternative unfolding patterns, indicating that this nearly completed SRP RNA transcript explores a folding landscape that branches into energetically closely degenerate, mutually exclusive folded states. Lastly, the F-X curve of full-length SRP RNA bound with RNAP (complex VI in Figure 2B) displays a single reproducible rip (red curve in Figure 2D), indicating the formation of its final functional stable long-hairpin structure. Furthermore, the F-X curve of complex VI is indistinguishable to that of an isolated full-length SRP RNA (Figure S1C-S1F), confirming that the entire RNA has emerged from the RNAP exit tunnel and adopted the native long-hairpin fold.

### Full-Length SRP RNA Readily Adopts a Stable Native Structure

Since the sequence at ~103-nt (complex V in Figure 2B) is the branching point between structures containing the non-native intermediate, and the native fold, we chose this complex to investigate the folding process in greater detail. Specifically, we wished to compare its behavior with that of the full-length SRP RNA (complex VI in Figure 2B) to assess the role of the hairpin tip-distal stem in the refolding process. To this end, we performed “force clamp” experiments to follow transitions between states accessible to the transcript at different forces starting at 15 pN and ending at 9.2 pN. As shown in Figure 3A, folded RNA states with different extents of base-pairing are characterized by distinct tether end-to-end extensions. Six different structures were observed in the refolding process of complex V (Figure 3A, 3C, and S2A). The first folding/unfolding event was observed around 15 pN (from U to I-1) and was identified as the formation of the non-native intermediate structure based on unfolding force and length change (red trace in Figure 3A). When the force is lowered to ~12.5 pN (yellow trace in Figure 3A), the nascent RNA continues to reduce its end-to-end extension, transitioning from I-1 to I-2 and to I-3, eventually attaining an intermediate fold, I-4. Formation of this quasi-stable structure (Figure 2E, bottom right) from the fully unfolded state (U) is predicted to involve ~83-nt with a decrease in contour length ΔC_L_ of 44.57 nm (= 83-nt × 0.59 nm/nt - 2 × 2.2 nm/RNA hairpin helix width), in good agreement with the contour length change measured between the unfolded state U and I-4, ΔC_L(U toI-4)_ = 43.8 nm (Figure 3C, peak-to-peak distance). When complex V is held at 10.5 pN (blue traces in Figure 3A), we observe a rapid fluctuation in tether extension, reflecting the interconversion of the RNA molecule between two structures differing in extension by ~7 nm. This extension change matches the expected transition length from quasi-stable to native-like structure (~ 16-nt × 0.59 nm/nt - 2.2 nm/hairpin helix width = 7.24 nm). This result is consistent with the aforementioned F-X curve (purple curves in Figure 2D), showing that the ~103-nt intermediate SRP RNA (complex V in Figure 2B) forms two competing structures at equilibrium—quasi-stable and native-like structures (Figure 2E, bottom).

**Figure 3.**
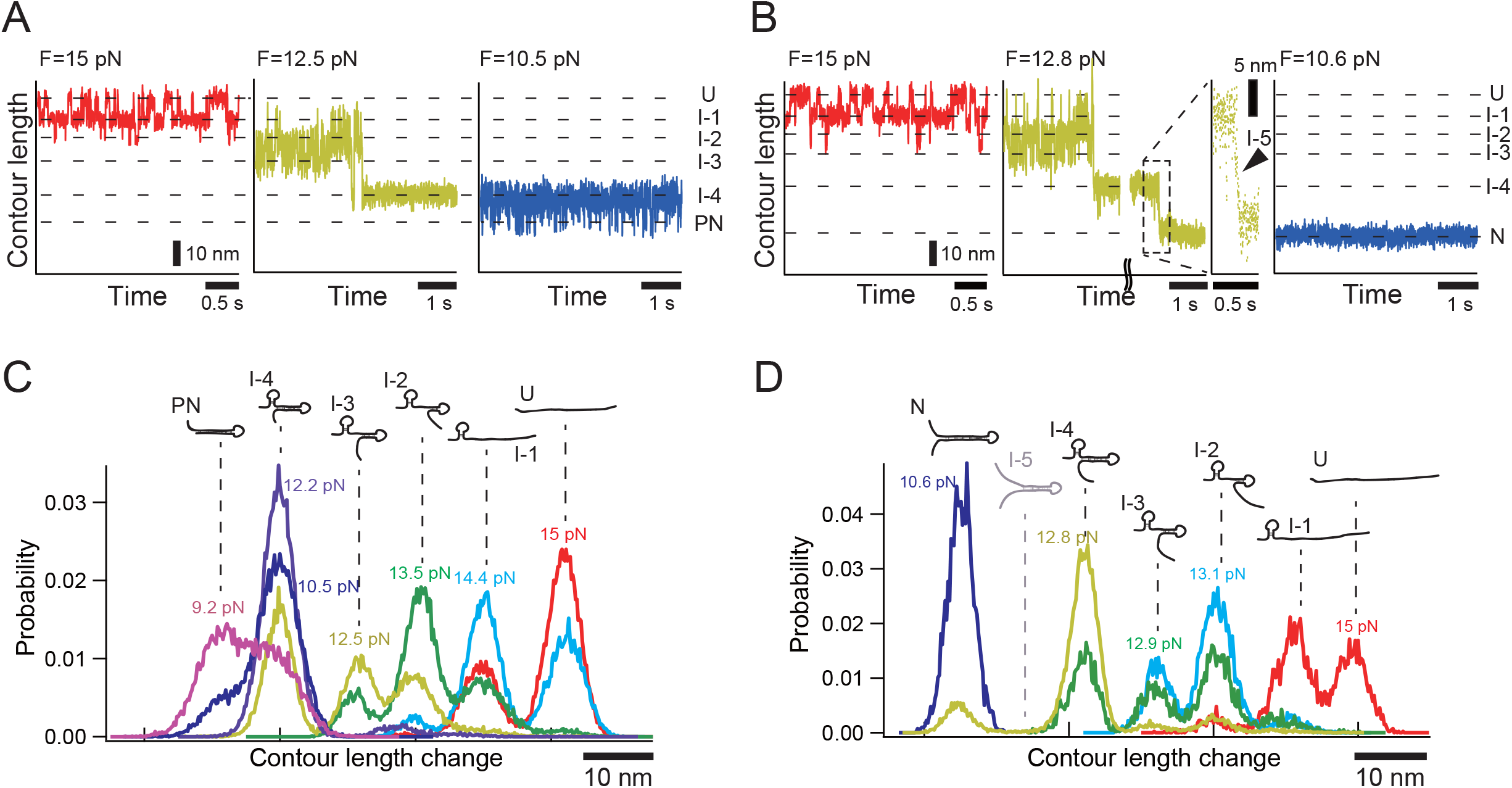
Comparison of Intermediate- and Full-SRP RNA State Determined by Refolding at Constant Force. Refolding trajectory of ~103-nt intermediate (A) and full (B) SRP RNA at different forces (3A and 3B). As force is reduced, non-native intermediate structure refolds (red). (A) The quasi-stable structure is highly populated around 12 pN (yellow). Two structures (quasi-stable and native-like structure) can exist at low force (blue). (B) The stable structure (N) forms at F = 12.8 pN. Enlargement of the trace at F = 12.8 pN (dashed box) reveals that additional state is visited immediately before forming stable structure (arrowhead). The structure (N) is stable over the observation and never unfolds at low forces. (C and D) Probability distribution of complete trajectories with ~103-nt intermediate SRP RNA (C) and full SRP RNA (D) at different forces, with scaled by the contour length. All trajectories to construct probability distributions (C and D) at each force are available in Figure S2.

Next, we repeated the same “force-clamp” experiment with complex VI (full-length SRP RNA) emerging out of RNAP. We identified six peaks representing six folding transitions (Figure 3B, 3D and S2B), where five of the states—U, I-1, I-2, I-3, and I-4—are identical to those characterized with complex V. However, we do not detect the rapid fluctuation (blue trace in Figure 3B) observed with complex V, corresponding to the transition between I-4 and a partially native state (PN, blue trace in Figure 3A) at low force (~10.5 pN). These observations indicate that the full-length SRP RNA predominantly adopts the final native fold, N, at the expense of the PN state. Close inspection of the trajectory at 12.8 pN shows that just before the formation of the stable structure (N), the RNA transiently passes through state I-5 (yellow trace in Figure 3B and inset of black box region). Thus, the full-length SRP RNA transitions through a total of seven structural folding states along the step-wise reduction of the forces applied in the mechanical refolding process.

The observation that the full length SRP RNA (complex VI) can transition from quasistable intermediate state I-4 to form the native structure under higher forces (12.8 pN), whereas the intermediate SRP RNA (103-nt, complex V) does not, can be rationalized thermodynamically. As calculated by Mfold, the free energy gain of the full-length SRP RNA when it adopts the native long-hairpin structure is twice as large as that of the non-native, quasi-stable intermediate (Figure 2E, bottom right): ΔG_native (full)_ = -250.6 kJ/mol < ΔG_quasi-stable (full)_ = -120.5 kJ/mol. In contrast, there is marginal free energy gain for the ~103-nt, truncated SRP RNA to transition from the quasi-stable structure to adopt the native-like fold (ΔG_quasi-stable (~103-nt)_ = -121.8 kJ/mol vs. ΔG_native-like (~103-nt)_ = -132.6 kJ/mol). Therefore, this analysis shows that the last ~9-bp on the hairpin tip-distal stem in SRP RNA contributes to form and stabilize the native structure, even though this portion of the hairpin stem (i.e. nucleotides 1-9 and 103-114) is known to be ultimately dispensable for SRP GTPase activity (Ataide et al., 2011; see last section below).

### Following SRP RNA Adopt Co-transcriptionally its Native Long-hairpin Fold

Next, we used the SRP RNA folding landscape obtained with truncated complexes at equilibrium to identify the structural transitions occurring in real time along the co-transcriptional folding pathway of the molecule. In these experiments, we used a similar geometry as Figure 2A to tether a single RNAP-RNA nascent chain complex, stalled after the T7A1 promoter at a short G-less sequence (right before the SRP RNA gene, beginning at a guanosine; see Methods), and held at a low constant tension (8.6 pN) to permit the folding of the RNA (Figure 4A). NTPs at various concentrations (10 μM to 1 mM) were then flowed in to re-start transcription. In these conditions, the emergence of the nascent RNA chain from RNAP can be followed in real-time as an increase in the tether end-to-end extension. An example of a co-transcriptional folding trajectory obtained in this manner is shown in Figure 4B (y-axis: extension in nm; x-axis: time in sec). Here a positive slope in the trajectory reflects the continuous growth (G) of RNA while a folding transition appears as a sudden decrease in tether extension (F). Regions of gradual negative slope seen in the trajectory are periods in which RNA growth and folding of the newly synthesized chain with pre-existing sequences occur concurrently (G+F), whereas horizontal segments (i.e. tether length invariant over time) correspond to transcriptional pauses of RNAP (P) on the template DNA.

**Figure 4.**
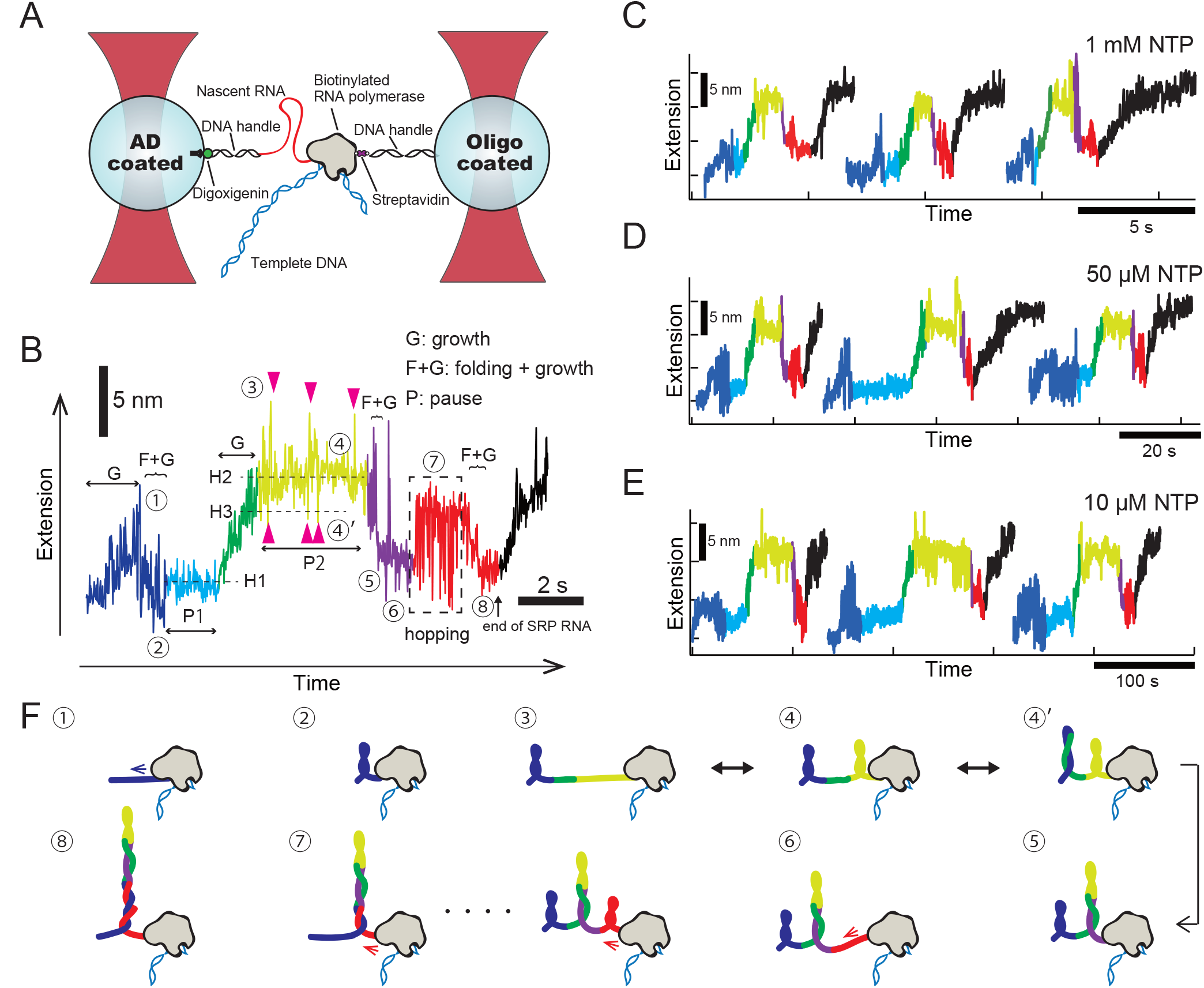
Real Time Transcription of SRP RNA. (A) Schematic of experimental setup to observe extension and folding of nascent RNA chain (not to scale). Two optical traps (red) were used to trap anti-digoxigenin (AD) and oligo coated beads. One DNA handle is attached to biotinylated RNAP via streptavidin and the other handle is hybridized to nascent RNA chain (see methods). (B) Representative trace of transcriptional extension of SRP RNA with 50 μM NTP (F = 8.6 pN). (C-E) Representative traces of transcriptional extension of SRP RNA with various NTP concentrations (F = 8.6 pN). Extension changes of nascent RNA chain were recorded at 800 Hz, averaged by decimation to 80 Hz (data for 1 mM and 50 μM NTP) and 16 Hz (data for 10 μM NTP). (F) Secondary structures of nascent SRP RNA during transcription are illustrated (not to scale), which are suggested by real time transcription traces.

### SRP RNA Displays a Remarkably Robust Co-Transcriptional Folding Pattern that Includes a Non-native Intermediate

Surprisingly, the co-transcriptional folding trajectory of SRP RNA is invariant and remains robust for up to three orders of magnitude change in NTP concentration (Figure 4C through 4E; N = 42 molecules; nucleotide K_d_ = 50 μM). Indeed, trajectories recorded at vastly different transcription rates can be re-scaled by their total duration and super-imposed onto one another. Consistently and reproducibly, these trajectories comprise three main features (Figure 4B): (1) a small hump (highlighted in dark blue); (2) two long plateaus (highlighted in light blue and yellow, respectively), where the latter one is embedded with spikes; and (3) a distinctive hopping in RNA extension (highlighted in red) before the completion of SRP RNA synthesis (Figure 4B, black arrow). Extensive control experiments were conducted to assign folding features in the trajectory to specific portions of the nascent RNA (see sections below; see also Methods for procedures to obtain the buffer-dependent conversion from extension in nm to molecular length in nucleotides, nt).

The first hump in the trajectory corresponds to the initial growth of the nascent chain until it reaches a length of 17-20-nt, followed by a subsequent folding transition in which it forms a small intermediate hairpin (H1) (Figure 4B and 5A). Note that after this initial folding event, the RNA extension drops approximately to the same level as the starting point of the trajectory. To verify the structure of intermediate hairpin (H1), we monitored how the co-transcriptional RNA folding pattern changes in the presence of an antisense DNA oligo complementary to the first 10 nt on the SRP RNA 5’-end (Antisense oligo-1 in Figure 5A and 5B). When we added the competing antisense oligo, this initial hump feature (Figure 5C, “F+G” region in the black trace)—present at all NTP concentrations tested and, therefore, co-transcriptionally *obligatory*—disappeared in the real-time folding trajectory (dark red traces in Figure 5C). Accordingly, the RNA tether extension becomes larger because the right-arm stretch of the H1 hairpin is now left as a single strand in the presence of antisense oligo-1 (Figure 5C, gray dashed lines around the second pause site P2).

**Figure 5.**
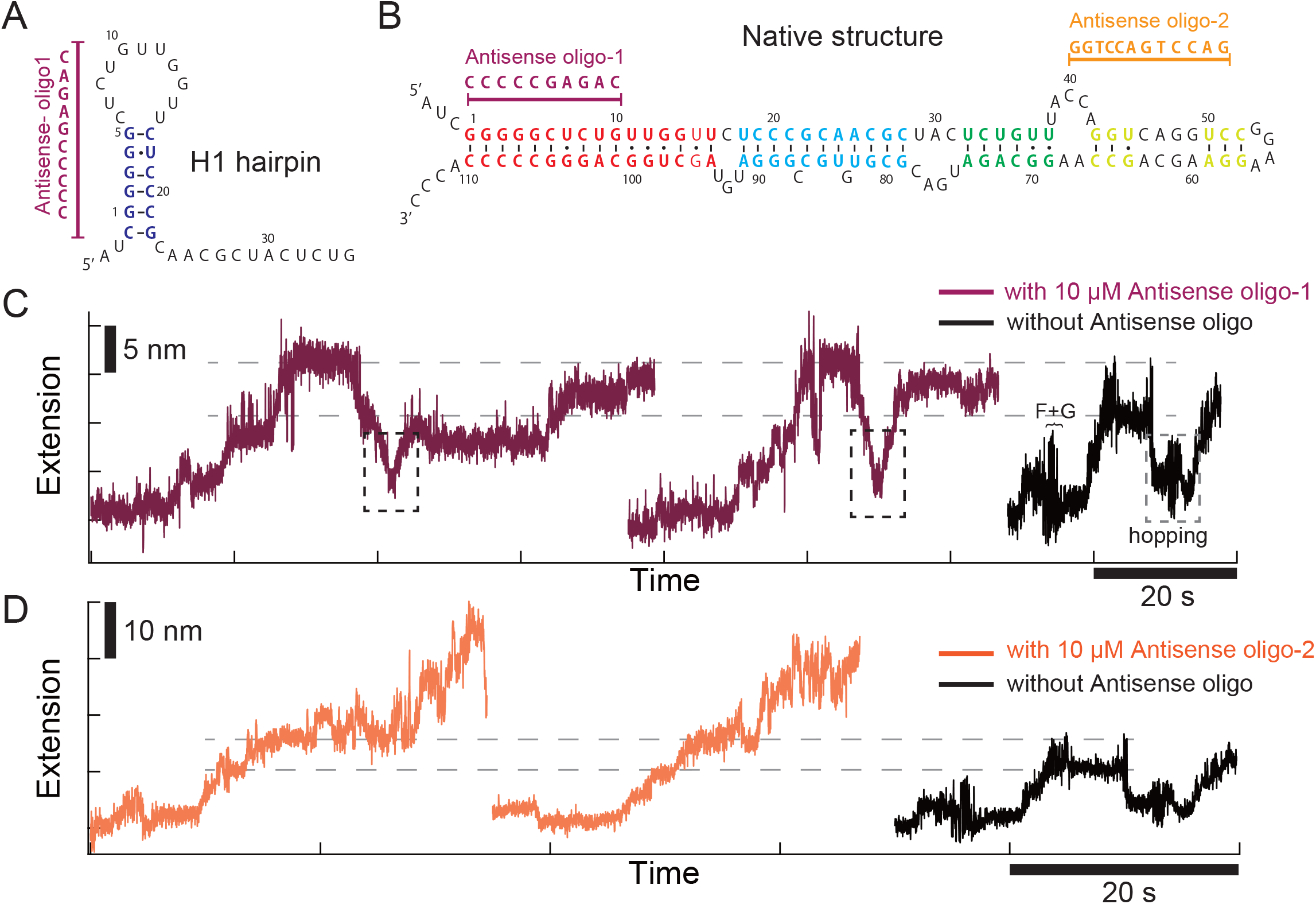
Real Time Transcription of SRP RNA with Antisense Oligo that Prevents the Nascent Transcript Folding. (A) Proposed secondary structure of non-native intermediate of SRP RNA (H1 hairpin in Figure 4B) and the sequence of antisense oligo. (B) Secondary structure of SRP RNA and antisense oligoes. The first antisense oligo (Antisense oligo-1) is designed to hybridize 5’ portion of specific intermediates (1-10-nt). The second antisense oligo (Antisense oligo-2) is designed to hybridize 5’ tip portion of native structure (40-50-nt). (C) Representative traces of transcription with the antisense oligo (Antisense oligo-1) with 50 μM NTP (dark red). For comparison, the standard transcriptional trace with 50 μM NTP (without antisense oligo) is shown (black). Gray-dashed boxes indicate the region of hopping dynamics observed in the absence of antisense oligo. Black-dashed boxes highlighted the disappearance of these hopping behavior in the presence of antisense oligo. “F+G” indicates the folding of non-native intermediate observed in the absence of antisense oligo. Dashed gray lines show the extension at second pause site with and without antisense oligo, respectively. (D) Representative traces of transcription with the antisense oligo (Antisense oligo-2) with 50 μM NTP (orange). For comparison, the standard transcriptional trace with 50 μM NTP (without antisense oligo) is shown (black). Dashed gray lines show the extension at second pause site with and without antisense oligo, respectively.

### SRP RNA Samples Alternative Structures at Pause Sites Along the Folding Trajectory

After the first intermediate hairpin (H1) forms, RNAP pauses (P1: light blue in Figure 4B), then resumes transcription for another ~55-nt (green in Figure 4B) and pauses again (P2: yellow in Figure 4B). Both pause durations increase when the NTP concentration is decreased (Figure S3A and S3B). To probe the origin and nature of these pauses, we conducted co-transcriptional folding experiments under various NTP-depletion conditions. When [UTP] is decreased to 10 μM, we observe selective lengthening in pause durations for both sites, P1 and P2 (Figure S4A, S4D and S4E); a low [GTP] also leads to pause lengthening at P2 (Figure S4B and S4E). In contrast, when ATP and CTP are depleted, no appreciable differences in both pause durations are observed compared to saturating conditions, i.e. 1 mM NTPs (Figure S4C). To accurately assign the locations of P1 and P2 in the SRP RNA sequence, we performed transcript elongation experiments while holding the RNAP-RNA complex under high tension (~22 pN) (Figure S5B though S5D). High force prevents RNA folding and yields a higher resolution transcription trajectory (Righini et al., 2018). We found that P1 is located around U36 (36.2 ± 2.5 nt into transcription), whereas P2 occurs around U82/U84—a known prominent pause site which has been shown to be suppressed by base-pairing compensating mutations U82A/A25U (Figure S5A), that preserve the long hairpin RNA structure (Wong et al., 2007). When we introduced these mutations, P2 was significantly shortened (Figure S5F and S5G), thus confirming our initial assignment. The G73U82G83 stretch in P2 matches perfectly with the consensus pause element G_−10_Y_−1_G^+1^ (where +1 position is the nucleotide to be next incorporated) (Vvedenskaya et al., 2014), thus leading to the strong G-pause detected at P2 (Figure S4B). As for P1, its U-rich sequence stretch U_34_G_35_U_36_U_37_U_38_—and similarly the U_82_G_83_U_84_ stretch at P2—resemble another pause-promoting element, U_−2_G_−1_, reported previously (Hein et al., 2011), hence resulting in long transcription pauses under limiting [UTP] (Figure S4A). The folding-free trajectories obtained at high force also reveal that the sequence-dependent P1 and P2 pauses do not require the formation of RNA secondary structure. Moreover, the duration of P1 and P2 pauses remained invariant to the addition of GreB (1 μM), a transcription factor that rescues RNAP from a back tracking pauses (Toulmé et al., 2000), indicating that both pause sites do not involve RNAP backtracking (Figure S3A and S3B).

Interestingly, during the second pause (P2), we detected upward and downward spikes corresponding to different discrete sizes in RNA extension (magenta arrowheads in Figure 4B). We propose that these features represent different partial folding states transiently accessible to the newly synthesized stretch of RNA. At this point, the nascent SRP RNA is ~70 nt long in total length and spends most of the time in a partial folded state, H2 (Figure 6B), which contains the aforementioned small intermediate non-native hairpin (H1) on the RNA 5’-end and, ~24 nt downstream, a second hairpin. Note that the second hairpin stem formed in H2 bears the essential binding site for FfH M-domain, i.e. the hairpin tip-proximal stem next to the tetra-loop (Figure 1B, blue-shaded region). This folded state (H2) can sometimes unfold (corresponding to the observed upward spikes) and quickly refold into a more compact but short-lived partially folded state, H3 (Figure 6C; corresponding to the downward spikes around P2 region in Figure 4B). The formation of the partially folded state H2 was confirmed by its inhibition with another antisense oligo (Antisense oligo-2 in Figure 5B): there is no drop seen in extension after the second pause site P2, showing that the downstream half of the hairpin tip-proximal stem (counting from the tetra-loop) is not base-paired and left as a single strand in the presence of antisense oligo-2 (Figure 5D, gray dashed lines around the second pause site P2). To confirm also the identity of the partially folded state H3, we introduced specific point mutations (U32C/U34C) aimed at preferentially stabilizing this putative state (Figure 6C; see also section below); the clear lengthening of residence time in the H3 state observed with these mutants (the downward spikes become well-resolved dwells, as seen in Figure 6E red traces), validate our assignment.

**Figure 6.**
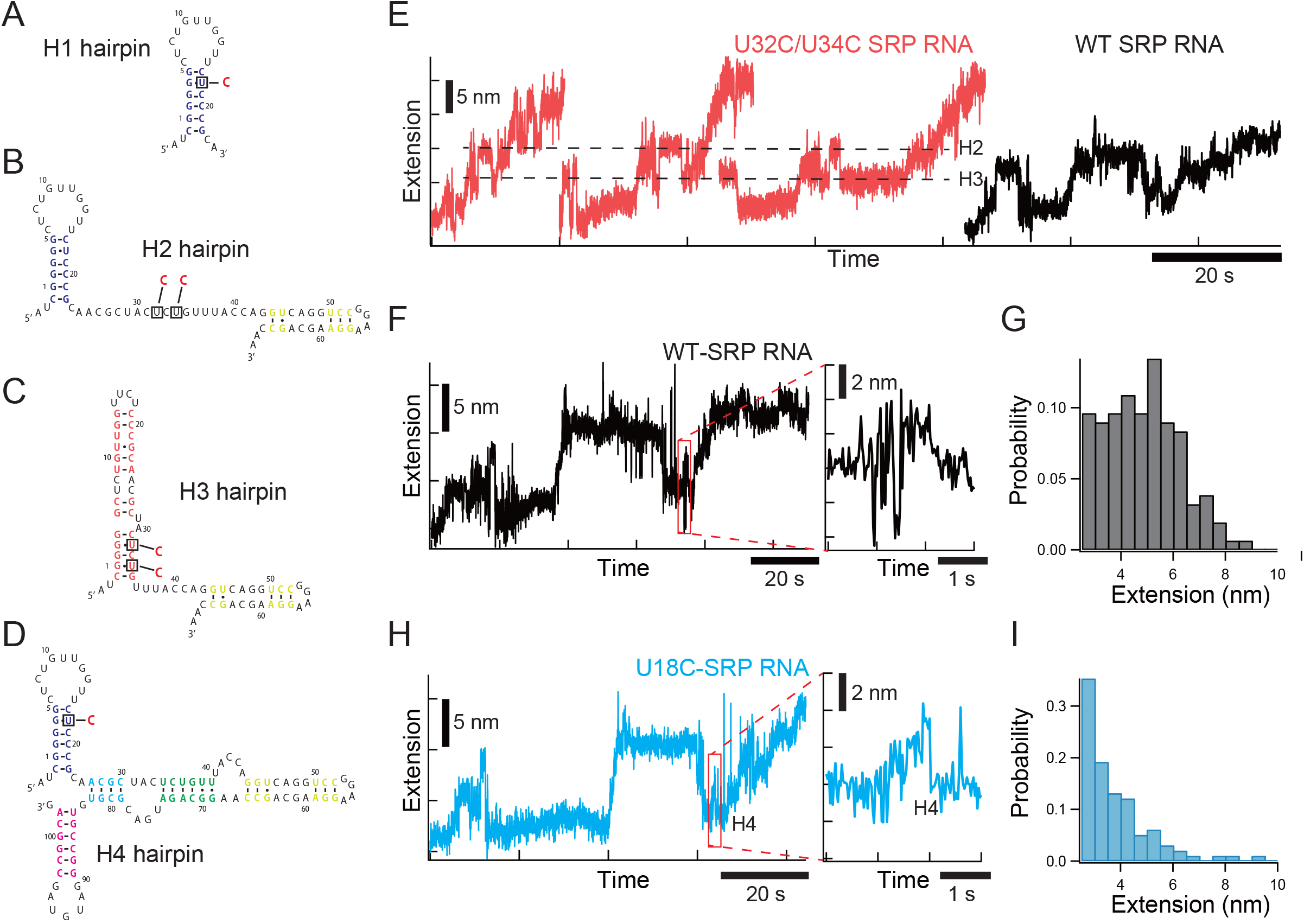
Real Time Transcription of Mutant SRP RNAs. (A-D) Expected secondary structures of H1 (A), H2 (B), H3 (C) and H4 (D) hairpins. U18 was mutated to stabilize the non-native intermediate structure (H1 hairpin). The free energy of H1 hairpin is increased from -29.3 kJ/mol (WT) to -41.8 kJ/mol (U18C) by the mutation as calculated by Mfold. U32 and U34 were mutated to stabilize H3 hairpin. The free energy of this structure (H3) is increased from -82.01 kJ/mol (WT) to -103.8 kJ/mol (U32C/U34C) by the mutations as calculated by Mfold. (E) Representative traces of transcriptional extension of U32/34C (bright red) and WT (black) SRP RNA with 50 μM NTP (F = 8.6 pN). (F and H) (Left) Representative traces of transcriptional extension of WT (black) and U18C (cyan) SRP RNA with 50 μM NTP (F = 8.6 pN). (Right) An enlarged trace (red box) shows structural transition from quasi-stable to native structure in WT SRP RNA (F). (G and I) Distribution of extensional change during structural switching (red box in F and H) of WT (G) and U18C (I) SRP RNA.

### SRP RNA Must Exit the Obligatory Non-productive Intermediate to Adopt the Final Native State

Eventually, RNAP exits this second pause (P2) and resumes transcription (purple in Figure 4B), adding nucleotides to the 3’-end of the ~70-nt long SRP RNA, which at this point spends most of its time in the partially folded state H2. The magnitude of the negative slope observed in this part of the RNA folding trajectory depends on the NTP concentration (Figure S3C), with lower concentrations leading to shallower slopes. This observation indicates that as the newly synthesized RNA emerges (in total 31-nt long), it readily base-pairs with the upstream complementary single-stranded region located between the two intermediate hairpins present in the partially folded state H2. Up to this point, ~60% of the final long-hairpin native fold of SRP RNA has formed. Next, distinctive hopping in RNA extension—i.e. recurring unfolding (large extension) and refolding (short extension)—appears toward the end of SRP RNA co-transcriptional folding trajectory (Figure 4B, region highlighted in red; Figure 6F and inset of red box region), reflecting RNA structural transitions among competing folding states. The average hopping amplitude is 4.5 ± 1.5 nm with a maximum of ~8 nm (Figure 6G), which corresponds to a total of ~21-nt under a constant tension of 8.6 pN; this change in RNA extension is consistent with the unfolding of the intermediate hairpin H1 (~7-bp stem + 5-nt loop). Also consistent, the first antisense oligo control experiment described earlier—designed to abolish the initial formation of intermediate hairpin H1—eliminates these hopping dynamics (black-dashed boxes in Figure 5C). These two features in the folding trajectory further establish that the hopping dynamics reflect the exiting of SRP RNA from the *obligatory*, yet *non-productive*, intermediate hairpin fold H1. As such, the hopping behavior represent the equilibrium between two major competing folding states of SRP RNA: the initial intermediate hairpin H1 plus ~60% of the final long-hairpin native fold, and the complete (~100%) long-hairpin native fold. Therefore, the low points in the hopping RNA extension correspond to the 5’-end stretch (~21 nt) of SRP RNA still sequestered in the H1 intermediate hairpin fold, while the high points in the hopping shows when the intermediate hairpin has unfolded. The hopping features are followed by a region displaying again negative slope, denoting concurrent RNA growth and folding. Thus, as the last segment of SRP RNA 3’-end emerges from the polymerase, it readily base-pairs with the 5’-stretch of SRP RNA previously sequestered in the *obligatory* intermediate hairpin fold H1 and transiently released through the hopping dynamics. Note that after this final folding event, the RNA extension drops approximately to the same level as that before the hopping—i.e., before the unfolding of hairpin intermediate H1 begins. These coincident RNA extensions suggest that similar amounts of base-pairing were involved before and after hopping—again, consistent with the interpretation that the SRP RNA 5’-end unfolds from the intermediate hairpin H1 and base pairs with the 3’-end stretch to adopt the final functional long-hairpin folding state. Note that it is only with this long-hairpin folding state that the hairpin tip *distal-stem*—i.e. the G-domain docking site essential for SRP’s protein targeting activity—is finally generated.

The very same distinctive hopping transition detected here in the co-transcriptional folding experiments was observed earlier in the “force-clamp” experiments, where the nearly completed (~103-nt long) SRP RNA stalled complex V—held at ~9-10 pN—rapidly switches between the *quasi-stable* intermediate state I-4 and the *native-like* long-hairpin fold, PN (blue trace in Figure 3A). However, the hopping transitions vanish when the full-length SRP RNA-RNAP (complex VI; blue trace in Figure 3B) is held under the same force. Therefore, the nonnative intermediate hairpin H1 is obligatorily and significantly sampled during the co-transcriptional folding trajectory. A cartoon diagram depicting the co-transcriptional folding pathway of SRP RNA resolved in our work is summarized in Figure 4F.

### SRP RNA Mutations Tune the Transition Between the Intermediate and its Final Native Fold

The co-transcriptional analysis above shows that an emerging SRP RNA undergoes multiple structural rearrangements between competing folding states (seen as transient spikes in RNA extension and as distinctive hopping dynamics; Figure 6F and inset of red box region). We then sought to shift the relative residence times of SRP RNA in these states by introducing two sets of point mutations: a U18C mutation (Figure 6A) that energetically favors the adoption of the intermediate hairpin H1, and mutations U32C/U34C (Figure 6C) that stabilize the rather shortlived partial folded state H3 detected during the transcriptional pause P2. In the case of U18C, the originally distinctive hopping dynamics in the co-transcriptional folding trajectory (Figure 4B, region highlighted in red; Figure 6F and inset of red box region) is attenuated (Figure 6H, inset of red box region, and 6I), confirming that the structural switching has shifted away from attaining the long-hairpin native fold. Replacing the hopping is a hump-like feature (similar to that seen with H1 hairpin formation), which indicates that the emerging, newly synthesized 3’-end single strand can sample a small hairpin fold H4 (Figure 6D and 6H; Figure 4F, cartoon #7) before resolving into the native long-hairpin fold. The mutation set U32C/U34C, besides similarly attenuating the hopping dynamics, converts the previously transient downward spikes into a clear dwell (Figure 6E, red traces). Therefore, the partially folded state H3 is more stably populated upon U32C/U34C mutations and hinders the transition into the functional long-hairpin fold. Interestingly, when we waited long enough after the entire SRP RNA sequence has emerged out of RNAP, we observed that both SRP RNA mutant constructs eventually adopt the functional long-hairpin native structure (i.e., their F-X curves exhibit the same unfolding pattern as that seen in the wild-type full-length SRP RNA; Figure S6A through S6C). This observation indicates that, thermodynamically, the structural folding equilibrium of these full-length SRP RNA mutants still lies toward the native folding state. Nonetheless, the co-transcriptional folding pathway data indicates that using minimal point mutations, we can trap kinetically the SRP RNA in a *non-functional* intermediate fold—which may function as an “off-switch” for its protein-targeting activity.

### Biasing SRP RNA Folding Equilibrium Strongly Affects Cell Viability

The real-time, co-transcriptionally-resolved folding features described here indicate that SRP RNA behaves like a molecular switch, converting between non-functional folding intermediates and the final long-hairpin native state. Although the intermediate and native folds of SRP RNA possess the hairpin tip-proximal stem essential for the recruitment of FfH and FtsY (Figure 1B), the hairpin tip-distal stem—responsible for the activation of SRP functionality—is critically missing from the former (Shen et al., 2013). Accordingly, it should be possible that biasing the equilibrium between alternative SRP RNA folds may influence its cellular activity. Hence, we sought to introduce mutations that bias the distribution of folding intermediates without preventing the adoption of final native state, and characterize their effect *in vivo*.

Since knockout SRP RNA strains are inviable (Doudna and Batey, 2004), we utilized a standard *E. coli* BL21 strain (non-T7 expression; NEB, C2530H), which produces native “switch-on” SRP RNA at endogenous levels, and transformed the bacteria with plasmids carrying intermediate-enhanced, potentially “switch-off” SRP RNA mutants. Three features of the plasmid constructs ensure the proper cloning and the precise synthesis of SRP RNA transcripts (Figure S7A; see Methods for plasmid sequences). First, an operon-embodying and tightly-repressible promoter, P_LtetO−1_ (Lutz et al, 1997), which only upon induction initiates efficient transcription. Hence, in a specialized *E. coli* strain that regularly expresses repressor genes integrated in the chromosome (*TetR* in DH5alphaZ1; Expressys), such a promoter prevents leaky transcription of potentially toxic SRP RNA mutants during cloning, permitting stable plasmid amplification. This operon-embodying—as opposed to operon-appending—promoter ensures the RNA transcript has a precise 5’-end. Second, a strong terminator, *rrnC* (Chen et al., 2013), immediately after the SRP RNA; and third, a second P_LtetO−1_ promoter located 54-nt downstream from the terminator. Binding of another RNAP to this second promoter serves as a roadblock to ensure that the first RNAP properly terminates on *rrnC*.

Based on the folding state populations characterized in our stalled RNAP-RNA complex experiments, we selected three SRP RNA transcripts whose folding equilibria are biased to various degrees towards the 5’-end-sequestering intermediate hairpin fold (quasi-stable structure in Figure 2E), inserted each sequence to the plasmid design described above, and tested the response of each transformed BL21 strain. These chosen transcripts were: (1) The full-length, wild-type SRP RNA (WT; 114-nt), which upon synthesis attains exclusively the native fold and does not populate the intermediate hairpin H1, thus serving as the “0%” reference. (2) A slightly shortened SRP RNA (SRP_100), where 14-nt on the 3’-end are deleted. This construct is seen to adopt the quasi-stable intermediate form (Figure 2E) 39.2% of the time, while the other 60.8% attains the activity-essential (G-domain docking site) hairpin tip-distal stem. (3) The same shortened SRP RNA but with a U18C mutation (U18C_100) to energetically stabilize hairpin H1 (Figure 6A). This truncation-plus-mutation combination further boosts the population of quasistable intermediate form to ~90% of occupancy: the majority of F-X curves show either two consecutive rips (19.9%; Figure S6E) or a series of small rips (69.9 %; Figure S6F), whereas a small fraction of F-X curves show single large rip (10.2 %; Figure S6D). We also made two control plasmids: (1) no SRP RNA sequence inserted between the promoter and the terminator (empty); and (2) the SRP_100 truncation variant but missing the second promoter downstream of the terminator so as to lessen the *precise* termination and generate read-through RNA transcripts (read-through).

As shown in Figure 7A, repression-free transcription of truncated SRP RNA variants— which enhance the intermediate fold by 40% or more—is lethal to *E. coli*, with few colonies surviving (bottom half, left and middle: SRP_100, U18C_100). In contrast, high-level synthesis of exogenous full-length wild-type SRP RNA (WT) has no impact on cell viability, yielding normal cell growth similar to that seen with the empty plasmid control (Figure 7A, top half, left and middle: WT, empty). We sequenced the few colonies surviving in the SRP RNA variant experiments, and found that these bacteria harbored plasmids with either mutations or truncations in the promoter or terminator (Figure S7B), which prevent the proper synthesis of the intended SRP RNA variants and likely alleviate the lethal impact of those plasmids. Indeed, the read-through control plasmid (Figure 7A, top half, right-most: read-through), though bearing the SRP_100 truncation variant, showed no deterioration on cell viability, hence confirming that it is the precise synthesis of the “intermediate-enhanced” SRP RNA transcripts that renders a lethal phenotype.

**Figure 7.**
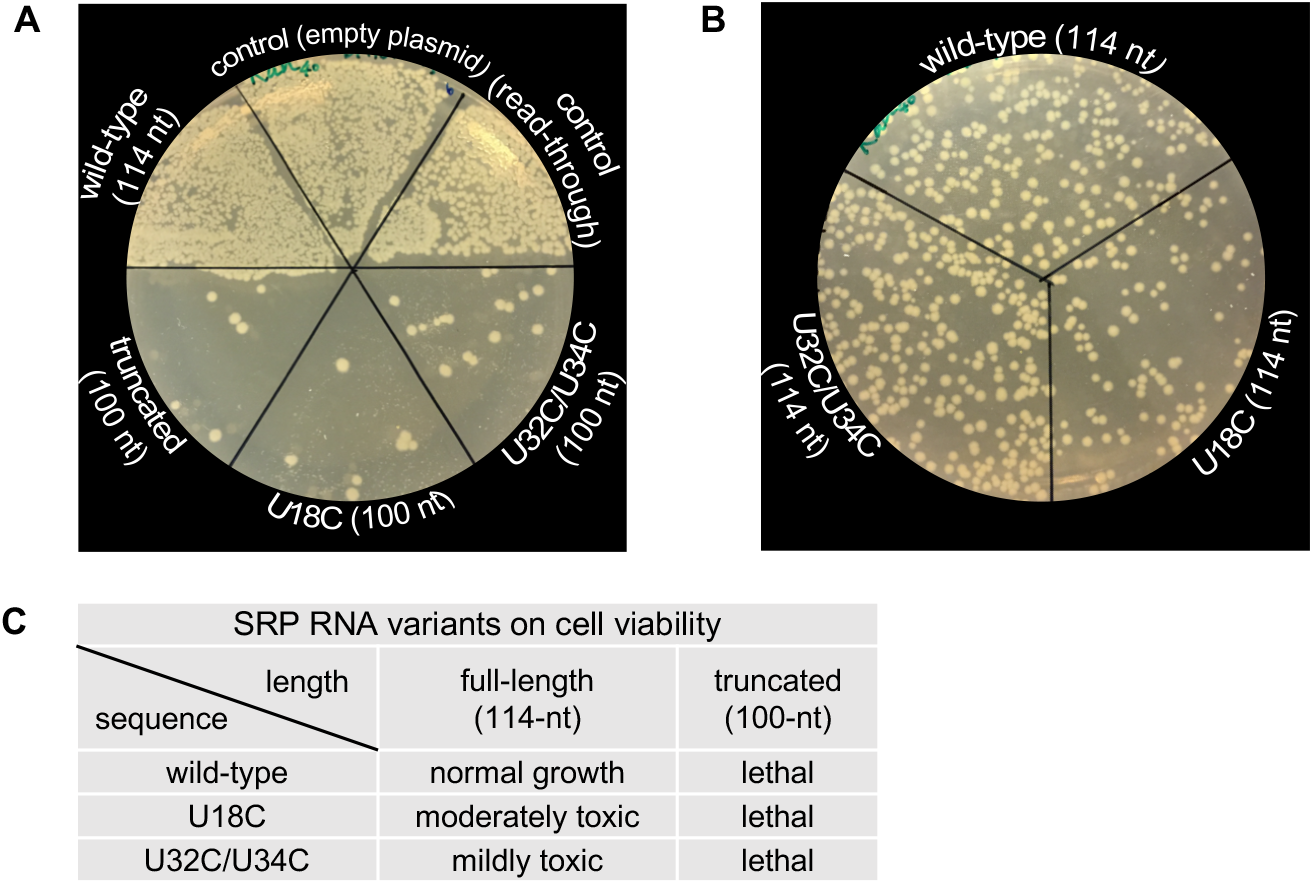
Impacts on cell viability from intermediate-stabilizing SRP RNA variants. (A) Plasmids carrying SRP RNA variants were transformed into *E. coli* BL21 strain, and plated on restriction LB plates (kanamycin, final conc. 40 μg/ml). While transcription of full-length wild-type SRP RNA (upper-half, left-most) shows normal cell growth as that of the control conditions (empty plasmid and transcript read-through construct: upper-half, middle and right-most), all the 3’-end truncated SRP RNA variants (SRP_100, U18C_100, U32C/U34C_100) are lethal. The surviving colonies were sequenced and identified to harbor modified plasmids (Figure S7B) that prevents the proper synthesis of the intended SRP RNA variants, thereby alleviating the lethal impact (see main text). (B) The full-length SRP RNA variants containing point mutations to stabilize specific co-transcriptional folding intermediates were also examined. While U18C_FL shows clear reduction in surviving colonies (lower-right), U32C/U34C_FL (lower-left) appears to have normal cell growth as that of wild-type_FL (upper-middle). The subtle differences in toxicity of full-length SRP RNA mutants were resolved by comparing their decrease in “post-relaxation” recovery growth rate to that of wild-type_FL (see main text, Methods, and Figure S7D-S7F). (C) Summary table on *in vivo* impact of all SRP RNA variants examined in this work.

The lethality of 3’-end-truncated SRP RNA variants indicates that the adoption of intermediate hairpin H1 by the 20 nt on the RNA 5’-end can sequester SRPs in an inactive/switch-off form, disrupting the formation of GTPase docking stem (nucleotides 10-15 and 96-101) and impairing SRP protein-targeting activity. We consistently observed a lethal phenotype even when another sequence mutation (U32C/U34C), which has no impact on the formation of intermediate H1 but only prolongs the lifetime of the transient intermediate hairpin H3 (Figure 6C), is added to the truncated SRP RNA variant (Figure 7A, bottom half, right: U32C/U34C_100). Interestingly, it has been reported that when both the 5’- and 3’-ends are shortened on the SRP RNA hairpin-distal stem (corresponding to nucleotides 1-9 and 102-114; black-dashed line in Figure 1B), SRP activity remains unaffected (Ataide et al., 2011), indicating that such blunt-ended truncation causes no folding interference with the formation of G-domain docking stem, hence preserving SRP’s functionality. Therefore, it is the presence of the 10-nt single-stranded 5’-stretch remaining in the asymmetric truncation construct SRP_100 that tips the folding equilibrium toward the nonfunctional intermediate state, with lethal consequences for the cell.

Next, we asked whether full-length SRP RNAs bearing point mutations that specifically alter its folding dynamics—i.e. that kinetically trap the RNA in *non-functional* intermediates along its co-transcriptional folding pathway without tilting the final equilibrium away from the native long-hairpin fold (Figure S6A through S6C) have a deleterious effect on cell viability. Again, we use U18C to stabilize the intermediate hairpin H1 (Figure 6A), and U32C/U34C to stabilize the intermediate hairpin H3 (Figure 6C). The U18C full-length construct exhibits a clear reduction in surviving colony counts—compared to that of wild-type—in the cell viability plate assay (Figure 7B); such decline can be further aggravated under stress condition (e.g. high salt growth medium, Figure S7C). The U32C/U34C full-length construct, however, appears to be indistinguishable to that of the wild-type. Since these plasmid constructs are not lethal, we resolved their relative cell toxicity by comparing the tendency of the transformed strains for plasmid loss under permissive—i.e. antibiotic-free—growth condition. Specifically, after relaxing the plasmid-carrying bacteria in selection-free over-night cultures (inoculating single colonies picked from restrictive plates after transformation), we challenge them with kanamycin, and measure the growth rate, to discern the fraction of the cellular population that has rejected the plasmid carrying both the antibiotic resistance and the SRP RNA variant (see Methods for detailed protocols derived from the naïve test in Chen et al., 2017). We found that both strains containing full-length mutant SRP RNA variants display a slowdown in recovery growth compared to that of the wild-type construct (colored arrows in Figure S7D through S7F). We interpret this result as evidence that the mutant RNA variants are more toxic that the wild-type construct. Of the two full-length mutants, the U18C construct—suggested by our singlemolecule experiments as a way to strongly bias the obligatory intermediate H1—leads to the slower recovery rate. The results of all SRP RNA variants tested on cell viability are summarized in Figure 7C.

## DISCUSSION

Using optical tweezers-based single-molecule experiments, we have shown that it is possible to uncover the complete folding trajectory of an essential RNA (SRP RNA) as it emerges on the surface of the *E. coli* RNA polymerase during active transcription. SRP RNA exhibits a remarkably robust co-transcriptional folding pathway, which is invariant to the rate of transcription. Along this pathway, we identified the initial formation of a non-native, obligatory intermediate H1, then the adoption of the hairpin tip-proximal stem, and followed by the sampling of alternative folding states. This non-native intermediate persists throughout most of the folding trajectory until the emergence of the SRP RNA 3’ end guides the formation of the hairpin tip-distal stem of the molecule, bringing about the final structural transition into the native-state long-hairpin fold. One thermodynamically possible mechanism to resolve hairpin H1 could involve a toehold-mediated RNA strand displacement (Zhang and Winfee, 2009) between the SRP RNA 3’-end single strand and the 5’-end arm of the intermediate hairpin H1. During toehold invasion, stretches of the 3’ single strand attempt to base pair with (i.e. toehold onto) the loop region of H1 hairpin until a strong enough hybridization is established—reducing the entropy cost for nucleotide bases on the two ends to search for alternative base-pairing—and eventually complete strand displacement. Such mechanism is being proposed in this current issue by Yu *et. al*. based on their SHAPE-seq data and molecular dynamics simulations. The large scale hopping dynamics observed in the co-transcriptional folding trajectory before the formation of the native form (Fig. 4B) may reflect those transient toeholding attempts between the two ends on SRP RNA.

The co-transcription folding trajectories reveal that a truncated SRP RNA missing the last 14 nt at the 3’-end primarily adopts a non-functional intermediate fold that lacks the essential G-domain docking site, hence *switching off* the protein-targeting activity of the SRP complex. This obligatory but non-productive intermediate can be exploited to structurally trap the molecule in an “off-switch” state, thereby enabling us to manipulate *E. coli*’s viability. Specifically, using this truncated SRP RNA (1-100 nt), we introduced mutations that alter the relative co-transcriptional residence time—and therefore the equilibrium—between the native state and alternative folds of the molecule (Figure 2E, bottom). Tilting the equilibrium away from the final native fold significantly reduces the number of SRP complexes competent for ribosome cargo transport and dramatically affects cell viability. Likewise, full-length SRP RNA mutants that do not prevent the final attainment of the native form but that kinetically affect the lifetimes of nonnative intermediates during the folding process, are seen to affect the ability of the cells to survive. The toxicity of these “kinetic” mutants is reflected in the increased degree to which these cells rid themselves of the plasmid-encoding SRP RNA variants. We speculate that, as both the endogenous and intermediate-enhanced SRP RNAs are synthesized in the cell, and as both transcripts possess the hairpin tip-proximal stem (Figure 1B), the two should compete to recruit the SRP proteins FfH and FtsY. Because the endogenous SRP RNA is known to be in excess of FfH by a factor of four in the untransformed cell (Jensen et al, 1994), and in our experiments the exogenous transcripts outnumber the endogenous ones, we expect the majority of FfH to be bound by the SRP RNA variants. In summary, we show that by modulating the interconversion between SRP RNA folding states revealed in the *in vitro* experiments, it is possible to harness SRP RNA as a molecular switch *in vivo*.

The discontinuous nature of transcription (interrupted by RNAP pauses) is thought to permit the RNAP-RNA nascent chain complex to interact with regulatory factors at different stages during transcription (Artsimovitch and Landick, 2000) altering the folding dynamics/pathway of the nascent RNA (Pan et al., 1999). Indeed, in the case of SRP RNA, we find that the second U-pause (P2; Figure 4B)—though not required for the formation of the hairpin tip-proximal stem—is retained in all co-transcriptional folding trajectories of full-length, wild-type SRP RNA. We speculate that it may instead play a role *in vivo*, perhaps providing a time-window wherein protein-RNA interactions crucial for the organization of the SRP complex take place.

While we found that the co-transcriptional folding pathway of SRP RNA is invariant to transcription rates, likely reflecting the simple long hairpin topology of the final native form of this molecule, other examples exist where the rate significantly reshapes the folding process (Lewicki et al., 1993). The methodology employed here should shed important insight into the coordination between synthesis and folding in those cases. Lastly, using SRP RNA as a model system, we envision that co-transcriptional modulation of the structural folding of other functional RNAs can be a generic way and direct portal to manipulate cellular processes. Conceivably, the SRP RNA variants characterized here could be exploited as toxins toward bacteria, serving as prototypes in the design of alternative antibiotics.

## STAR⋆METHODS

### SUPPLEMENTAL INFORMATION

Supplemental information includes Extended Experimental Procedures, 7 figures, 1 table, and Supplemental References and can be found with this article online at *****

## ACKNOWLEDGEMENT

We thank L. Alexander for critical reading of the manuscript and all members of Bustamante laboratory for critical discussion. We also acknowledge J. Lucks, A. Yu and E. Strobel (Northwestern University) for helpful discussion. The pZE21 vector used in the cell experiments is a kind gift from A. Flamholz (Savage lab at UC Berkeley). This work was partly supported by the MEXT/JSPS Grants in Aid for Scientific Research JP15K17889, and the US Department of Energy Office of Basic Energy Sciences Nanomachine Program under Contract DE-AC02-05CH11231.

## AUTHOR CONTRIBUTIONS

Conceptualization, S.F. and S.Y.; Methodology, S.F. and S.Y.; Investigation, S.F., S.Y., and Y.K.; Writing - Original Draft, S.F., S.Y., R.G., and C.B.; Writing — Review & Editing, S.F., S.Y., and C.B.; Funding Acquisition, C.B.; Resources, R.G., M.S., S.F., and S.Y.; Supervision, C.B.

